# KaScape: A sequencing-based method for global characterization of protein-DNA binding affinity

**DOI:** 10.1101/2023.06.19.545523

**Authors:** Hong Chen, Yongping Xu, Jianshi Jin, Xiao-dong Su

## Abstract

It is difficult to exhaustively screen all possible DNA binding sequences for a given transcription factor (TF). Here, we develop a method named “KaScape”, by which, TFs bind to all possible DNA sequences in the same DNA pool where the DNA sequences are prepared by randomized oligo synthesis and the random length can be adjusted to, e.g., 4, 5, 6, or 7, etc. After separating the bound from unbound double-strand DNA, their sequences are determined by next-generation sequencing. To demonstrate the relative binding affinities of all possible DNA sequences determined by KaScape, we develop a three-dimensional KaScape viewing software based on a K-mer graph. We apply KaScape to 12 plant TF family *At*WRKY proteins and find that all *At*WRKY proteins bind to the core sequence GAC with similar profiles. KaScape not only can detect binding sequences that are consistent with the consensus W-box “TTGAC(C/T)”, but also other sequences with weak affinity. KaScape provides a high-throughput, easy-to-operate, sensitive, and exhaustive method to quantitatively characterize the relative binding strength of a TF to all possible binding sequences, allowing us to comprehensively characterize the specificity and affinity landscape of transcription factors, particularly for moderate and low affinity binding sites.

**Highlights:**

- A general method, KaScape, using NGS (next-generation sequencing) with a series of random dsDNA for exhaustive characterization of protein-DNA binding.
- A K-mer-based analysis and display software tool KGViewer developed to analyze the relative affinity landscape of KaScape results.

## 1. Introduction

The interaction between transcription factors (TFs) and their specific transcription factor binding sites (TFBSs) ^1^ is crucial for TFs to regulate gene expression ^2^. The current consensus is that TFs need to search along the double-stranded DNA (dsDNA) before they bind to their TFBSs, and there are weak or moderate affinity binding sites besides specific high-affinity TFBSs. In fact, some moderate or low-affinity TFBSs may also be important for gene regulations^3^. Thus, a comprehensive understanding of the interaction between TFs and DNA is essential which requires high-throughput (HTP) analytical methods. Recently, due to the development of HTP, high-accuracy, and scalable next-generation sequencing (NGS) technologies, the NGS has been revolutionizing the study of TFBSs, enabling researchers to investigate the binding events of TFs on genome-wide scales ^4-6^. NGS-based methods such as chromatin immunoprecipitation followed by sequencing (ChIP-seq)^7^ and systematic evolution of ligands by exponential enrichment (SELEX)^8^ have been developed and widely used to identify and map TFBSs across the genomes. ChIP-seq, as an *in vivo* method, may exhibit non-negligible false positive outcome since the cross-linking used in ChIP-seq may make proteins become covalently trapped on non-specific chromosomal DNA and the antibody used in ChIP-seq may non-specifically bound to an untargeted TF ^9^. The *in vitro* HTP SELEX technique yields data with a lower degree of noise. However, SELEX usually can only identify enriched high-affinity TFBSs since it removes the low-affinity TFBSs during measurement cycles ^10^.

To overcome the limitations of existing methods based on NGS, and exhaustively identify TFBSs of a specific TF including all high-, and low-affinity TFBSs, we developed a method, named KaScape, which determined the relative binding affinities of all possible DNA sequences to the interested TFs (for example, the *At*WRKY family DNA binding domains). In KaScape, a library containing all possible 4-, 5-, 6-, or 7-bases of dsDNA sequences of which the composition was determined by NGS, was firstly mixed with each of the TFs of *At*WRKY family; secondly, the bound TF-DNA complexes were separated from the mixture; thirdly, the DNA sequences in the separated TF-DNA complexes pool were determined by NGS; finally, the relative binding affinities of all possible dsDNA sequences were calculated based on both the proportion of each DNA sequence in the separated TF-DNA complexes pool and the original DNA library. Furthermore, to demonstrate and analyze the relative binding affinities of all possible DNA sequences determined by KaScape using a K-mer graph in three dimensions, we have developed a program suite KGViewer.

## 2. Materials and methods

### 2.1. Random dsDNA preparation

We prepared a random dsDNA pool by extending another DNA strand on random single-strand DNAs (ssDNAs) with randomized 4, 5, 6, or, 7 bases in the middle, and fixed sequences on both flanking ends using a primer (see Table S1 Complementary ssDNA). The ssDNAs were synthesized by Integrated DNA Technologies, USA (see Table S1 Random ssDNA, n represents the random base length). The ssDNAs were mixed with the primer (synthesized by Sangon, China) and EasyTaq PCR SuperMix (the reagent concentrations are listed in Table S2). Then the mixture was incubated using a polymerase-chain-reaction (PCR) thermo-cycler with a program shown in Table S3. The dsDNAs were purified by gel filtration with superdex75 (GE Healthcare, USA). The purified dsDNAs are called random dsDNA pool. We note that the length of the dsDNA is about 30 base pairs.

### 2.2. Protein preparation

The N- or C-terminal DNA binding domains (DBD) of *Arabidopsis* WRKY family proteins (*At*WRKY1, *At*WRKY2, *At*WRKY3, *At*WRKY4, *At*WRKY32, and *At*WRKY33) used in this study were prepared using *E. coli* BL21 as previously reported ^11^. Briefly, the codon-optimized genes of the DBDs were constructed into the pET21b vector with C-terminal His-tag. The constructed vectors were subsequently transformed into the *E. coli* strain BL21 (DE3). The transformed bacteria were induced by adding isopropyl β-D-1-thiogalacto-pyranoside to a final concentration of 0.5 mM and then were grown overnight at 18°C to express the DBDs. To purify the DBDs, the bacterial cells were collected and resuspended in buffer A (25 mM HEPES, pH 7.0, 1.0 M NaCl), followed by sonication and centrifugation. Afterwards, the DBDs in the supernatant were purified using a Ni-chelating column and a size-exclusive chromatography (Superdex 75, GE Healthcare, USA); the DBDs were finally eluted in buffer C (25 mM HEPES, pH 7.0, 100 mM NaCl). The purified DBDs were stored at - 80°C after flash freezing by liquid nitrogen.

### 2.3. KaScape procedures

The procedure of KaScape includes the following five steps (Fig. 1):

**Fig. 1.**
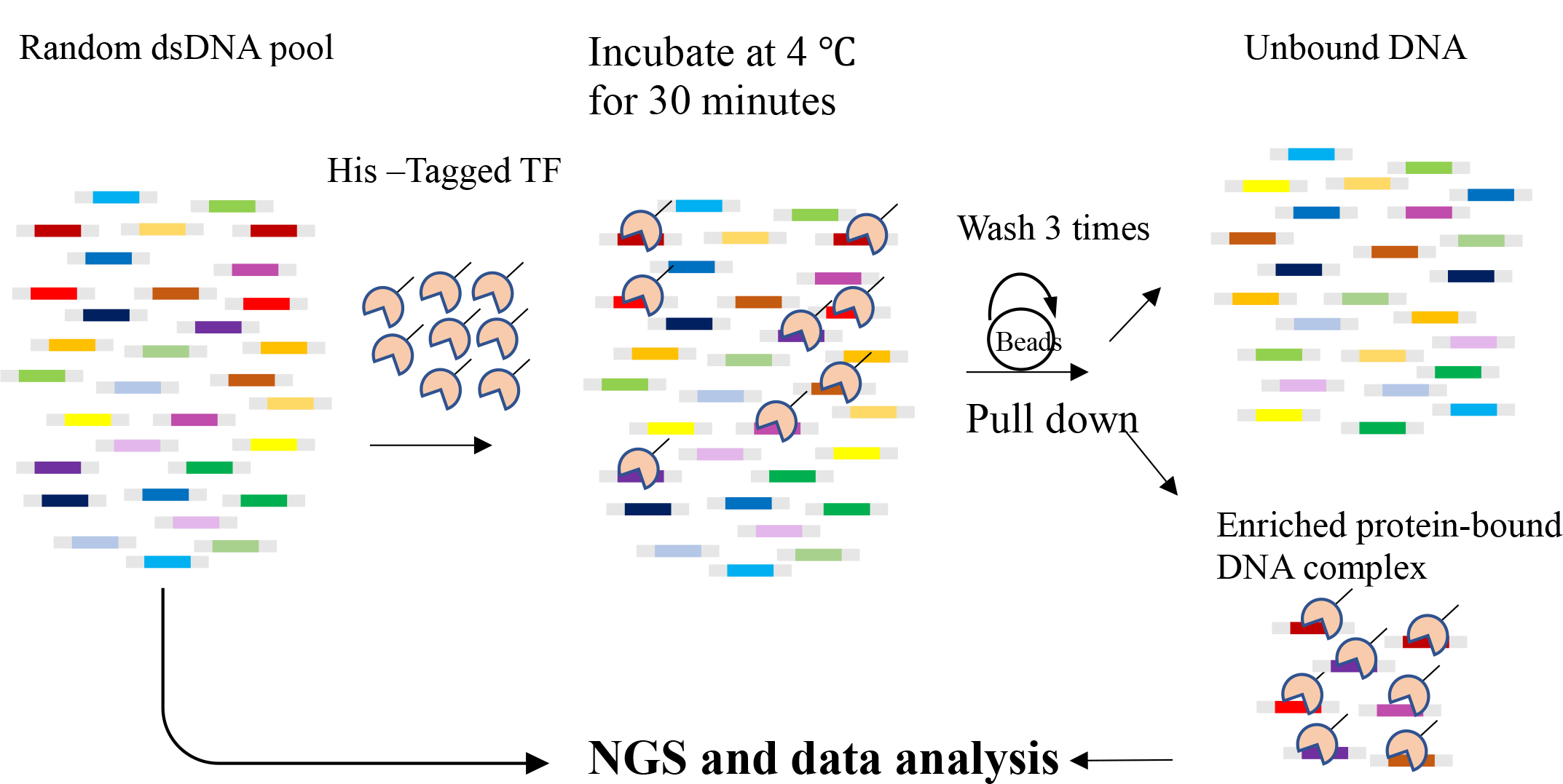
Schematic description of the KaScape process, not to scale. The random dsDNA pool of n=4,5,6,7 as colored rectangular bars above, and His-tagged TF DBD were prepared. The pooled dsDNA is about 30 base pairs with flanking sequences (Table S1). The 2 × 10^−11^ mol protein and 10^−10^ mol dsDNA were mixed into 2 mL buffer. Then the TF DBD and dsDNA were incubated for 30 minutes. Next, the magnetic His-tag purification beads were added and rotated for an hour, and then the system was washed and rotated 3 times. After that the dsDNA and TF DBD complex were separated with free unbound dsDNA. Finally the random dsDNA pool and bound dsDNA were extended and constructed to the dsDNA library separately for next-generation sequencing.

#### 2.3.1 Mixing protein with dsDNA

For each of the *At*WRKY proteins, 2 × 10^−11^ mol DBDs and 10^−10^ mol random dsDNA pool (about 1 µg) were mixed with buffer C to the final volume of 2 mL with a EP tube, and were incubated on ice for 30 minutes.

#### 2.3.2 Separating bound and unbound dsDNA

First, the magnetic His-tag purification beads (BeaverBeads™ IDA-Nickel, Beaver for life science, China) were balanced with buffer C by following the product instruction. Second, 10 µL of the balanced magnetic beads were added to the protein-dsDNA mixture. Third, the mixture was gently rotated (approximately 10 rpm) for one hour at 4°C using HS-3 vertical mixer (SCIENTZ, China). Forth, after the magnetic beads were clearly separated from the mixture on a magnetic stand, the supernatant was removed slowly with a pipette. Fifth, the magnetic beads were suspended in 1 mL buffer C by rotating at 10 rpm for one hour using HS-3 vertical mixer (SCIENTZ, China). Sixth, after the magnetic beads were clearly separated from the mixture on a magnetic stand, the supernatant was removed slowly with a pipette. The fifth and sixth steps were repeated three times in total. Seventh, the magnetic beads were suspended in 50 µL of 500 mM imidazole by pipetting and incubated for 2 minutes at room temperature. Eighth, after the magnetic beads were clearly separated from the mixture on a magnetic stand, the supernatant containing protein-DNA complexes was transformed into a new EP tube. The bound dsDNA was purified from the transformed supernatant using the Oligo Clean & Concentrator Kits (ZYMO Research, USA) by following the kit instructions.

#### 2.3.3 Extending the dsDNA

The random dsDNA pool and purified bound dsDNAs were respectively extended to 75 bp for random sequence type in which the random base length (n) is 4 (76-79 bp for n equals 5-7) by PCR using an extension primer (Table S1 extension primer; synthesized by Sangon, China). The purified bound dsDNAs were mixed with the extension primer and EasyTaq PCR SuperMix (the reagent concentrations are listed in Table S4). Then the mixture was incubated using a polymerase-chain-reaction (PCR) thermo-cycler with a program shown in Table S5. The extended dsDNAs were purified using DNA Clean & Concentrator Kits (ZYMO Research, USA) by following the kit instructions.

#### 2.3.4 Library preparation and sequencing

Customized Illumina sequencing adaptors (Table S6) were ligated to the extended random dsDNA pool and extended bound dsDNAs respectively; mixing solutions for ligation were prepared as Table S7 and were incubated at 25°C for 20 minutes. The ligated dsDNAs were purified using AMPure XP beads (Beckman Coulter, USA) by following the product instruction with a ratio of 1:1.5 between the ligated dsDNAs and AMPure XP beads. For each ligated random dsDNA pool and bound dsDNAs after purification, different Illumina indexes were added using PCR. The PCR solutions were prepared as in Table S8 and the PCR program is shown in Table S9. Finally, the indexed libraries were purified using AMPure XP beads (Beckman Coulter, USA) by following the product instruction with a ratio of 1:1 between the library and AMPure XP beads for two times. The purified libraries were sequenced using Illumina NovaSeq PE150 platform.

#### 2.3.5 Sequencing data analysis

The random sequences between the designed fixed sequences were extracted from read 1 and read 2. If the extracted random sequences in read 1 and read 2 in the same pair were not reverse complementary, the pair of reads was discarded. For the remained reads of the sequencing results obtained from the random dsDNA pool, the number of reads for each type (*S*_*i*_) of random sequences was counted as 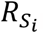 For the remained reads of the sequencing results obtained from the bound dsDNAs, the number of reads for each type (*S*_*i*_) of random sequences was counted as 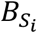. Then the proportion of the sequence *S*_*i*_ in the random dsDNA pool was calculated as 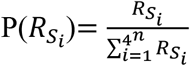; whereas, the proportion of the sequence *S*_*i*_ in the bound dsDNAs was calculated as 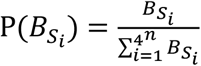. Finally, the relative intensity which represents the relative affinity of the sequence *S*_*i*_ among all possible DNA sequences was calculated as 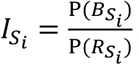. The above data analysis was performed using custom code written in Python 3.6. The modules re, os, sys, datetime, collections, gc, pandas, numpy, math, matplotlib, pickle, xlwt were used.

### 2.4. The K-mer graph

The theory of K-mer graphs was described in previous studies^12-15^. The 1-mer, 2-mer, 3-mer, and 4-mer graphs were shown in Fig S1.

### 2.5. KGViewer visualization software

The KGViewer visualization software (Fig. 3) was written in Python 2.7. The MayaVi scientific visualizing tool^16^ and QtGui python graphical user Interface display tool were used.

## 3. Results and discussions

### 3.1 The development of KaScape

To determine the binding ability of all kinds of dsDNA to a protein exactly under the same experimental condition, measuring the proportion of each dsDNA sequence type bound to a protein in the same liquid mixture is an ideal solution if the initial dsDNA distribution is even. To achieve this goal, a dsDNA pool containing all kinds of dsDNA sequences is required which can be made by random ssDNA synthesis. Since the binding sites of most single-domain transcription factors are short, (for example, the length of TFBS bound by WRKY is around 6 bps), we designed a random dsDNA pool using 4,5,6, and 7 random bases respectively. In an ideal scenario, each bound dsDNA molecule should be bound by only one transcription factor, which should be precisely located on the random base region. The more a specific type of DNA is measured, the stronger the binding signal for that type of dsDNA will be. Based on the aforementioned guidelines, we developed a new method called KaScape (Fig. 1). Due to certain biases during random ssDNA synthesis manufacturing, the random sequence distribution may be uneven. We thus sequenced the random pool and the bound dsDNAs, trying to define the relative intensity (see 2.3.5) that is the bound dsDNAs distribution divided by the random pool distribution to represent the relative binding affinity.

In a KaScape experiment, we first prepared a DNA pool including all possible sequences with a certain random base length n (n may be 4, 5, 6, or 7) using randomized ssDNA synthesis and dsDNA generation (see 2.1). The length of the oligonucleotide used for binding experiments is around 30 nucleotides (nt) and the Tm value of the Complementary ssDNA (see Table S1) is estimated at about 60°C. To evaluate the evenness of the random dsDNA pool, the pool was sequenced (see below). We calculated the random dsDNA pool distribution (Fig 2**a**) through the sequencing results (2.3.5). Figure 2**a** showed a slight bias and most differences in abundance among sequences were < 6. The read depth bias is due to the base utilization in ssDNA synthesis with the utilization order T>A>C>G (Fig. 2**d**). Next, we incubated the randomized dsDNA pool with a transcriptional factor DBD (see 2.2 and 2.3.1)). Our preliminary experiments show that a slight excess of DNA is more appropriate. Too much initial DNA will cause the final data dominated by high-content sequences of the initial DNA library due to the uneven input DNA distribution. Whereas if there is too little DNA, the specific DNA cannot be highlighted among the non-specific DNA sequences due to the experimental noise. The amount of protein used is determined by the amount of DNA at the molar ratio 1:5. As DNA may get lost in every step, the initial amount of DNA is about 1 µg. The longer the length of the randomized region, the more DNA amount is needed. 1 µg is enough for initial DNA with a random base length as short as 4-7 nt. In order to remove non-specific binding as much as possible, we washed the DNA-protein complex three times (see 2.3.2). we found that one time, two times, and three times of washing in KaScape did not significantly change the results (Fig. S2**b**); the washing step was the only essential step that might change the protein-DNA binding equilibrium for different sequences in our design.

**Fig. 2.**
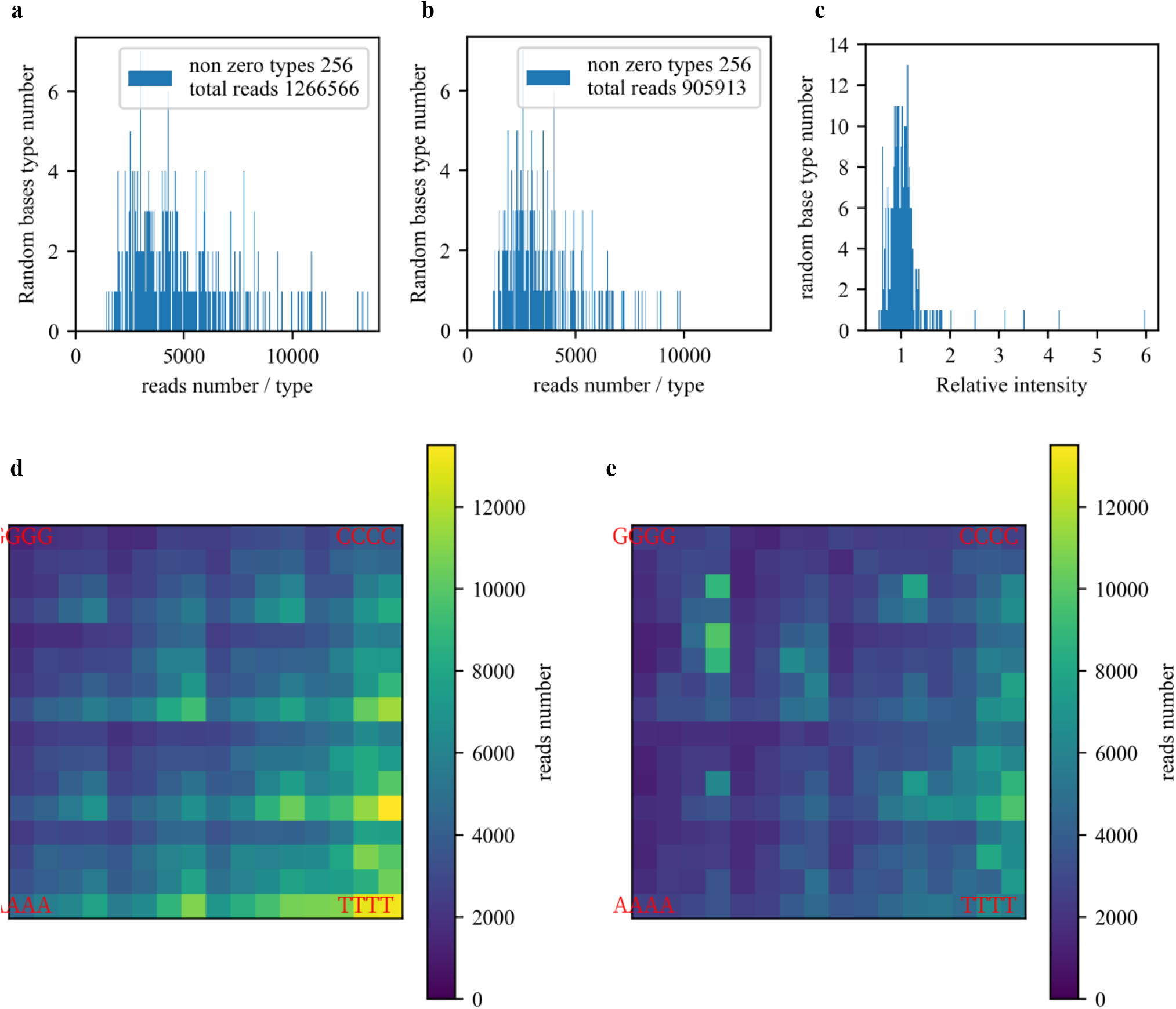
KaScape raw sequencing data. The random base number in the DNA sequence is 4. (**a**) The random dsDNA pool distribution. Each bar represents random base type number for a certain range of reads’ count numbers. (**b**) The bound DNA distribution. (**c**) The DNA relative intensity distribution. (**d**) The random dsDNA pool’s read depth landscape in K-mer graph. (**e**) The bound dsDNAs’ read depth landscape in K-mer graph.

We then isolated the bound DNA by pull-down (see 2.3.2). Due to the uneven random dsDNA pool, both the random dsDNA pool and the bound dsDNAs libraries were need to be prepared. The length of the dsDNA is too short to build a library for NGS. Thus before preparing the dsDNA library, we extended the random dsDNA and bound dsDNAs into DNA fragments of more than 70 bp respectively (see 2.3.3). There are 15 nt overlap between the Extension primer (see Table S1) and the dsDNA to ensure extension efficiency. The Taq DNA Polymerase we used in the extension step has terminal transferase activity resulting in the addition of a single nucleotide (adenosine) at the 3’ end of the extension product, which is convenient for subsequent library construction. The extended dsDNA is readily ligated to sequencing adaptors without having to conduct the “end repair” and “adenylation” steps required in many commercial kits for NGS. Compared with commercial kits, this method saves time and cost. Finally, the random dsDNA pool and bound dsDNA libraries were constructed and sequenced by next-generation sequencing (see 2.3.4). To evaluate the binding ability of different kinds of dsDNA, we analyzed (2.3.5) the distribution of the bound dsDNAs (Fig. 2**b**) and compared the bound dsDNAs with the random dsDNA pool. Figure 2b shows the distribution is narrower and there is a left shift in the peak compared to the random dsDNA pool distribution (Fig. 2**a**). Several dsDNA sequence types are highly enriched in the bound dsDNA (compare Fig. 2**d** and Fig. 2**e**). However, there is also a similar pattern in both the random dsDNA pool and bound dsDNA read depth landscape which is due to nonspecific binding (compare Fig. 2**d** and Fig. 2**e**). In order to characterize the relative affinity, we defined the relative intensity (2.3.5). Based on the relative intensity values, it appears that most of the sequences bound with the protein through non-specific binding, as their values were either close to or less than 1. This suggests that the protein may not have a strong affinity for these sequences. (Fig. 2**c**). However, several sequences exhibited a value significantly greater than 1, indicating that they were strongly bound sequences. (Fig. 2**c**). We confirmed that the relative intensity values of all sequences were highly reproduced in repeated experiments (Fig. S2**a**).

Since the next-generation sequencing depth is critical, we assessed the sequencing depths for KaScape by simulation. We adopted the sequencing depth requirement calculation from the paper ^17^, Fig. S3. If the random base length of dsDNA is 4, the library size is 256, assuming the ddG range between dsDNA and protein is between 0.5 and 5 kcal/mol, we need more than 100000 reads to achieve close to 100% accuracy. For the paired-end 150 sequencing strategy (PE150), one needs at least 30M. To investigate the sequencing depth requirement from experimental data, we calculated the correlation coefficient of relative intensity distribution between the experimentally derived data and several randomly down-sampling simulated data (see Fig. S4). The result is similar to the adopted calculation method (compare Fig. S3 and Fig. S4).

Therefore, KaScape method robustly characterized the relative binding landscape of all possible TFBS at the same time.

### 3.2. KGViewer, a K-mer-based 3-D visualization software

To study dsDNA interaction with a DBD protein (e.g., protein recognition and binding to a certain DNA sequence), measuring a value (e.g., affinity or relative intensity defined in this paper see 2.3.5) for all possible DNA sequences with thermodynamic equilibrium conditions exhaustively is the best way we can do in vitro (3.1). However, the global comparison of a measured value for all possible sequences (when the length is > 2) is difficult. To directly compare a value among all possible sequences, we developed a software, named KGViewer (see 2.5 and Fig. 3). KGViewer can visualize the given values of all possible sequences using the height and the color of bars plotted on a K-mer-based graph (Fig. S1).

**Fig. 3.**
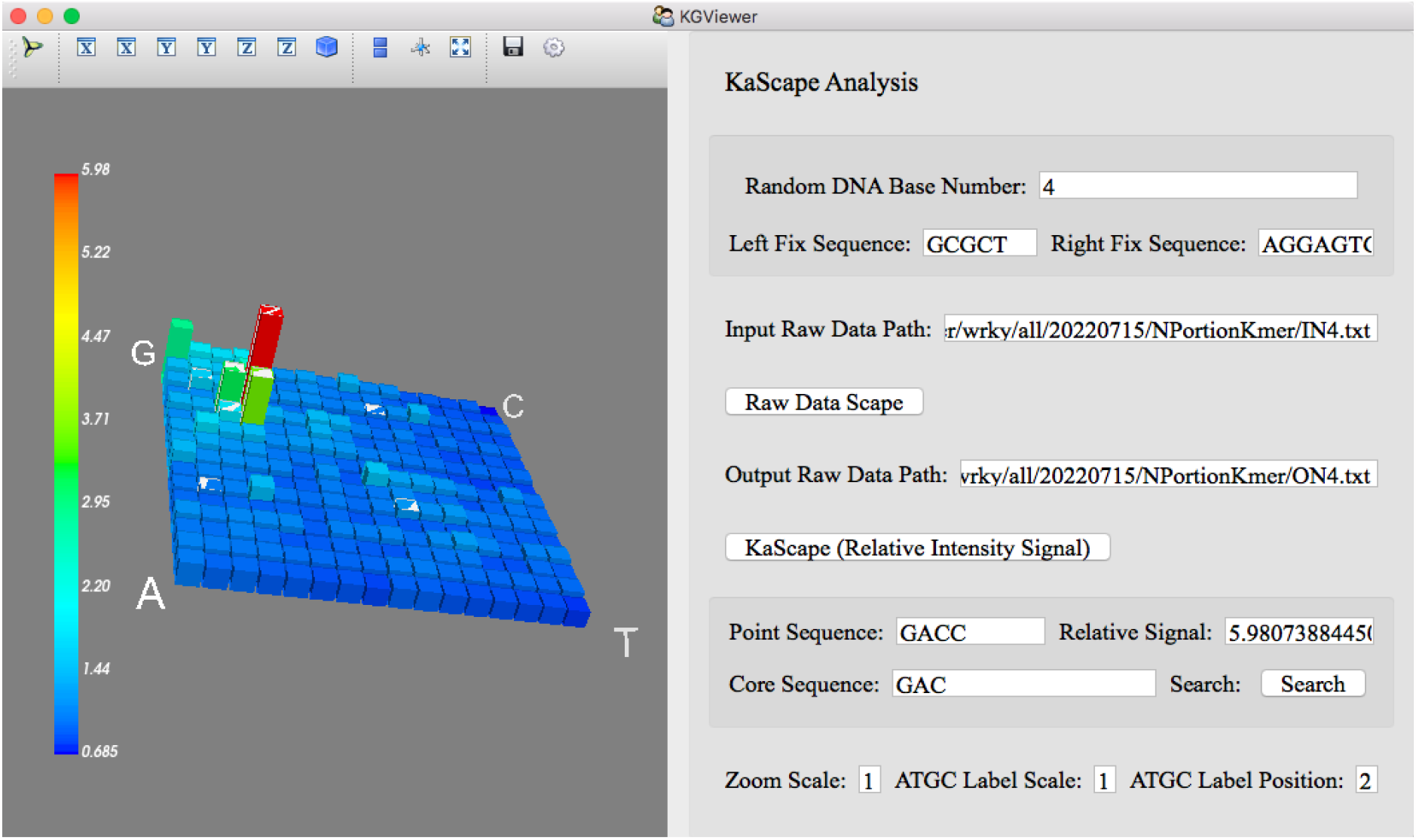
KaScape 3-D software KGViewer. The random dsDNA pool distribution and the relative intensity landscape can be shown in the left panel in K-mer graph. The sequence and signal value can be seen once the color bar is clicked (for example, GACC and 5.98). The highlighted sequence (see the shining bar) contained the core sequence searched by the user (Here NGAC or GACN are highlighted by searching the core sequence GAC, N represents one of the four bases).

The KGViewer has the following functions. The size of the K-mer-based graph (2, 3, 4, 5, 6, and 7 bases) can be adjusted by setting the parameter “Random DNA Base Number”. The random dsDNA pool distribution landscape can be conveniently visualized in a K-mer graph on the left panel by providing the input raw data path (the path to the random dsDNA pool distribution landscape file) and clicking on the “Raw Data Scape” button. After entering the output raw data path (the path to the bound dsDNA distribution landscape file) and clicking on the “KaScape (Relative Intensity Signal)” button, the left panel will be changed to show the relative intensity landscape. To view the value and sequence information of a bar in the landscape, the user can click on the bar, which will display the relevant information in the text box labeled “Point Sequence” and “Relative Signal”. For instance, clicking on a bar may show information such as the sequence ‘GACC’ and a value of 5.98. To visualize all sequences containing a core sequence (e.g. ‘GAC’) that one is interested in, a search function was provided to highlight them (e.g. ‘NGAC’ or ‘GACN’ that contain GAC was highlighted, N represents one of four bases). The landscape size, the G, C, A, and T labels’ size, and location can be scaled at the last row of the right panel.

This software KGViewer with flexible functions can visualize and directly compare a value of all possible sequences with a fixed length in a single plot, which is useful for investigating the distribution of a value in a K-mer-based space.

### 3.3. The binding affinity landscape for WRKY proteins

We applied KaScape to the WRKY family proteins (Fig. S5). In order to evaluate the overall binding affinity of the N terminal domain of *Arabidopsis* WRKY1 (*At*WRKY1N), we constructed a series of KaScape experiments using random base lengths (n) of 4, 5, 6, or 7. To facilitate the interpretation of binding affinity data obtained from the KaScape experiments, the K-mer graph is employed to arrange the relative intensities in a clear and concise manner. (Fig. 4). The relative intensity exhibits a proportional increase as the color shifts from purple to yellow. The four relative intensity landscape maps exhibit similar patterns. To assess the consistency across the series of KaScape experiments, we derived the K-mer relative intensity landscape map from the (K+1)-mer relative intensity landscape map, Fig. S6. By comparing Fig. S6**a**, Fig. S6**b**, and Fig. S6**c** with Fig. 4**a**, Fig. 4**b**, and Fig. 4**c** correspondingly, the patterns are similar respectively. The correlation coefficient of Fig. S6**a** and Fig. 4**a**, Fig. S6**b** and Fig. 4**b**, Fig. S6**c** and Fig. 4**c** are 0.292, 0.865, 0.74 respectively. The results of the correlation coefficient analysis suggest that the KaScape experiments conducted for random base lengths of 5, 6, and 7 are highly consistent. The signals in random base length 4 KaScape experiments’ relative intensity map is cleaner (compare Fig. 4**a** and Fig. S6**a**). In Fig. 4, A small region with strong signals is consistently observed in the upper left portion of all the K-mer graphs. The sequences within this region correspond to GACN, GACNN, GACNNN, and GACNNNN (N represents one of the four nucleotides (G, C, A, or T)), where n equals 4, 5, 6, or 7. When the n equals 4, the relative intensity order in the upper left signal region is as follows: GACC > GACT > GACG > GACA (Fig. 4**a**). This tendency remained when n becomes larger. As an illustration, when n equals 5, the relative intensity order is generally GACCN > GACTN > GACGN > GACAN.

**Fig. 4.**
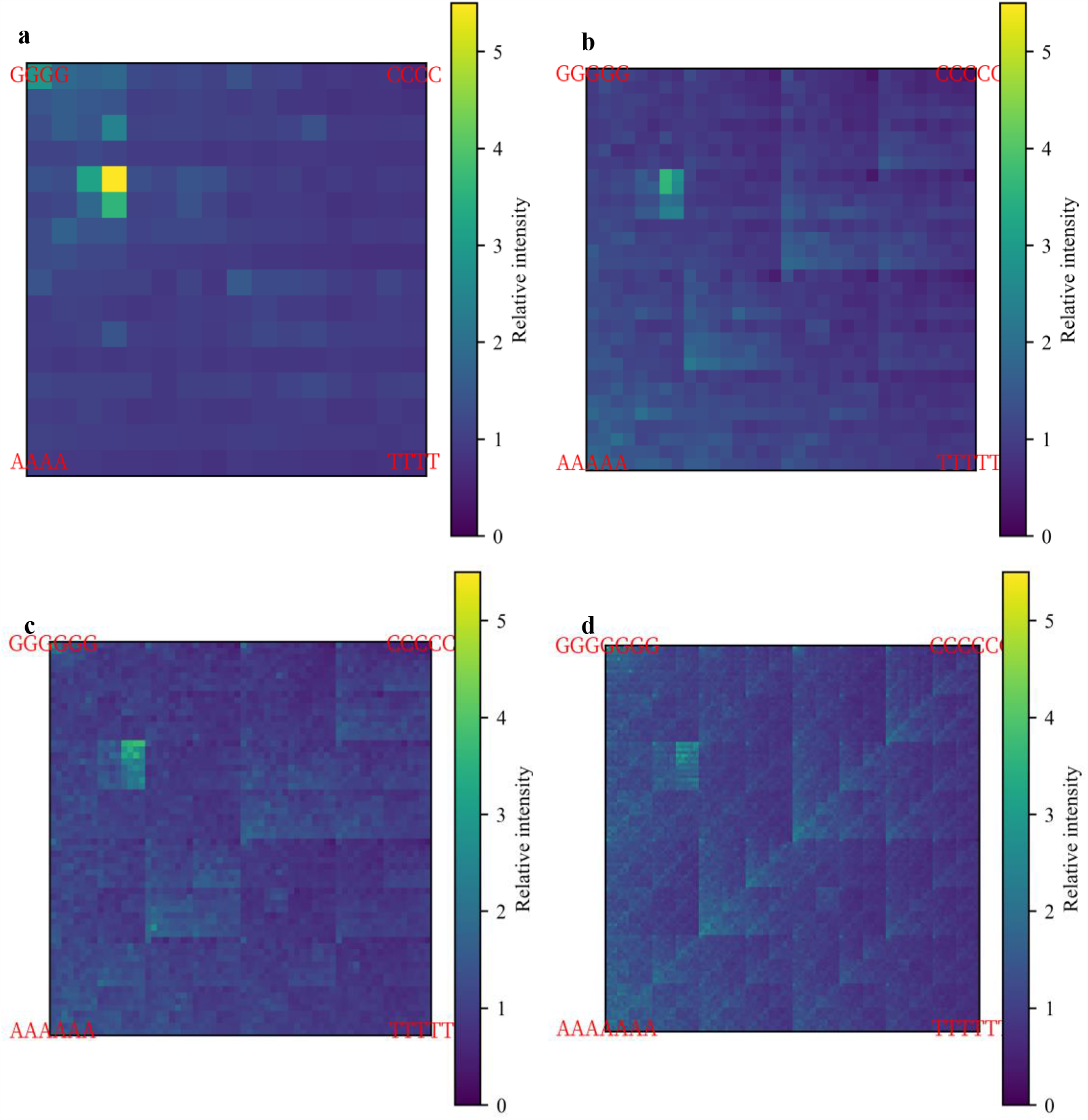
*At*WRKY1N KaScape relative intensity landscape on a K-mer graph for a series of KaScape experiments with random base length dsDNA sequences. (**a**) The random base length of the sequences used in the KaScape experiment is 4. (**b**) The random base length of the sequences used in the KaScape experiment is 5. (**c**) The random base length of the sequences used in the KaScape experiment is 6. (**d**) The random base length of the sequences used in the KaScape experiment is 7.

To confirm the KaScape results, we did EMSA experiments for 8 sequences that contain GACCN or NGACC (N represents G, C, A, or T), Fig. S7. *At*WRKY1N is capable of binding all those sequences except for the sequence which contains CGACC. The relative intensity of CGACC is 0.75, whose relative intensity is the lowest of the 8 sequence types used in the EMSA experiment. And the relative intensity of the other 7 sequence types is higher than or equal to 1.0. The EMSA results are consistent with the KaScape results. To gain insight into the binding specificity of the sequences with the highest affinities, we utilized PWM sequence logos to analyze the KaScape experimental data. Fig. S8 shows the sequence logos of the highest relative intensity sequences from each KaScape experiment of different random base lengths. The specific sequences are “GAC(C/T)”, “GAC(C/T)”, “GACC” and “GGTC” (or GACC in reverse complementary) when n equals 4, 5, 6, and 7 respectively. They are consistent in the series of KaScape experiments. The recognition of W-box “TTGAC(C/T)” ^18^ by the WRKY domain has been reported, however, subsequent studies have revealed that the predominant binding contribution arises from a shorter core sequence “GAC” (or “GTC” in reverse complementary) for the WRKY family in general ^11,19^ (Fig. S5). The specific sequence we generated here is part of the W-box and contains the core sequence GAC (or GTC in reverse complementary). The core sequence information is already contained in the random base length 4 KaScape experiment. Besides those bound sequences containing core sequence GAC, there are non-canonical motifs for WRKY family proteins to bind, such as ‘CAACA’ which *Th*WRKY4 can specifically bind ^20^. The relative intensity of ‘CAACA’ in the random base length 5 KaScape experiment is 1.5506. The relative intensity is larger than 1 which means ‘CAACA’ can enrich the binding of the transcription factor compared to the non-specific binding sequences. Thus the KaScape results can not only give us information about the core sequence of the canonical W-box motif but also provide non-canonical information ^21^. As it provides the relative affinity landscape for all possible sequences, KaScape can identify not only high-affinity binding sequences but also low-affinity binding sequences^22^ that are specific to the transcription factors being studied.

In order to see whether the core sequence is conserved in WRKY family proteins, we did 12 KaScape experiments for the N terminal and C terminal DNA binding domains of *At*WRKY1, *At*WRKY2, *At*WRKY3, *At*WRKY4, *At*WRKY32, and *At*WRKY33 (see the multiple sequence alignments in Fig. S5). As the core sequence information is already contained in random base length 4 KaScape experiment, Fig. S8. The random base length of random dsDNA for the KaScape experiments used here is 4. Fig. 5 shows the relative intensity landscape maps for *At*WRKY family proteins. The maps are similar. And most correlation coefficients of relative intensity between each *At*WRKY family protein pair are more than 0.9 (Fig. S9**b**). This indicates the core sequence is conserved for *At*WRKY family proteins. The correlation coefficients of relative intensity between *At*WRKY3C and the other 11 *At*WRKY family proteins are less than 0.8, as depicted in Fig S9. Additionally, the relative intensity landscape map of *At*WRKY3C displays the lowest signals when compared to other *At*WRKY family proteins, as shown in Fig 5. These observations suggest that *At*WRKY3C may exhibit weaker binding specificity and binding affinity than the other tested *At*WRKY family proteins.

**Fig. 5.**
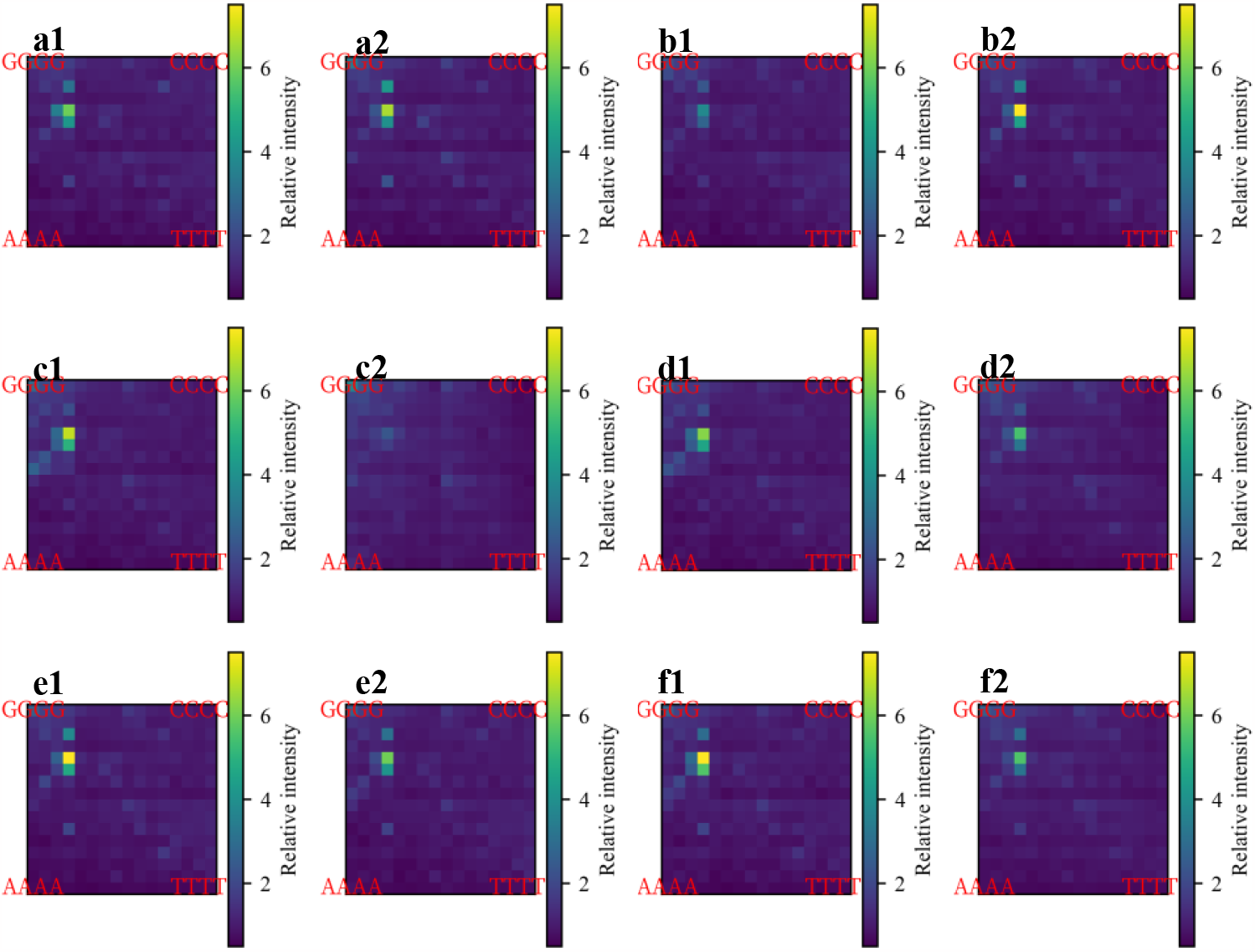
KaScape relative intensity landscape map for proteins in *At*WRKY family. (**a1**) *At*WRKY1N, (**a2**) *At*WRKY1C, (**b1**) *At*WRKY2N, (**b2**) *At*WRKY2C, (**c1**) *At*WRKY3N, (**c2**) *At*WRKY3C, (**d1**) *At*WRKY4N, (**d2**) *At*WRKY4C, (**e1**) *At*WRKY32N, (**e2**) *At*WRKY32C, (**f1**) *At*WRKY33N, (**f2**) *At*WRKY33C. ‘N’ represents the N terminal domain of WRKY, while ‘C’ represents the C terminal domain of WRKY, the random base length of all subfigures is 4.

## Author contributions

X.-D.S. conceived and supervised the entire project. Y.X. mainly performed the experiments, H.C. did all analyses and programming, X.-D.S., H.C. and J.J. discussed the data and wrote the manuscript.

## Data availability

The KGViewer software and the sequencing data analysis scripts are distributed freely and are available through the GitHub repository (https://github.com/NinYuan/KaScape.git).

